# GIT2 is dispensable for normal learning and memory function due to a predominant brain GIT2 splice variant that evades GIT/PIX complexes

**DOI:** 10.1101/538223

**Authors:** Krisztian Toth, Amanda C. Martyn, Natalia Bastrikova, Woojoo Kim, Ramona M. Rodriguiz, Umer Ahmed, Robert Schmalzigaug, Serena M. Dudek, William C. Wetsel, Richard T. Premont

**Author notes:** Corresponding author at: Department of Medicine, Case Western Reserve University, Cleveland, OH 44106 *E-mail address:. These investigators contributed equally to these studies (Krisztian Toth, Amanda C. Martyn). Current address: Campbell University School of Pharmacy, Buies Creek, NC 27506. Current address: North Carolina School of Science and Math, Durham, NC 27705. Current address: Harrington Discovery Institute, University Hospitals Medical Center, Cleveland OH 44106 and Institute for Transformative Molecular Medicine, Department of Medicine, Case Western Reserve University School of Medicine, Cleveland, OH 44106.

## Abstract

G protein-coupled receptor kinase-interacting protein 2 (GIT2) and GIT1 are highly similar, sharing the same domain structure and many binding partners. The most important GIT partners are the p21-activated protein kinase-interacting exchange factor (PIX) proteins, since through homomeric and heteromeric interactions, GIT and PIX proteins form oligomeric GIT/PIX complexes. Oligomeric GIT/PIX complexes function both as regulators of small GTP-binding proteins and as scaffolds for signalling molecules, including p21-activated protein kinases (PAKs). Deficits in learning and memory have been demonstrated in GIT1 knockout mice, and it has been assumed that GIT2 also would affect learning and memory. Unexpectedly, we find that GIT2-deficient mice respond normally in multiple tests of learning and memory, and have normal hippocampal long-term potentiation. Further, we find no evidence that GIT2 regulates ADHD-like phenotypes. To investigate why GIT2 and GIT1 differ so markedly in the brain, we identified the major isoform of GIT2 in the brain as a previously uncharacterized splice variant, GIT2(ΔBCE). This variant cannot dimerize or form oligomeric complexes with PIX proteins, and is thus incapable of regulating PAK in synapses, compared to oligomeric GIT1/PIX complexes. Because localized activation of PAK in synapses is required for structural plasticity underlying cognitive performance, loss of monomeric GIT2(ΔBCE) in the brain does not influence these responses.

## Introduction

The GRK-interacting (GIT) proteins, GIT1 and GIT2, are signalling scaffold proteins (Zhou, Li et al. 2016). The GIT proteins function as direct signal mediators through their ADP-ribosylation factor (Arf) GTPase-activating protein domain, which inactivates Arf family small GTP-binding proteins (Premont, Claing et al. 1998, Vitale, Patton et al. 2000). The GIT proteins also serve as subunits within an oligomeric scaffolding complex, formed together with p21-activated kinase-interacting guanine nucleotide exchange factor (PIX) proteins (Zhou, Li et al. 2016). GIT protein dimers formed through a coiled-coil self-association interact with PIX protein coiled-coil trimers (Schlenker and Rittinger 2009) to form very high-molecular weight oligomeric GIT/PIX complexes (Premont, Perry et al. 2004, Totaro, Tavano et al. 2012) that may consist of two or more of these presumed pentameric units. The tight linkage of GIT and PIX proteins is evident in the profound loss of PIX proteins in the brain of GIT1-deficient mice (Won, Mah et al. 2011) or in immune cells from GIT2-deficient mice (Hao, He et al. 2015), or of GIT2 in immune cells from a-PIX-deficient mice (Missy, Hu et al. 2008).

The two GIT proteins, GIT1 and GIT2, have been implicated in learning and memory function. This was based initially on the identification of the GIT/PIX signalling pathway members a-PIX (Kutsche, Yntema et al. 2000) and PAK3 (Allen, Gleeson et al. 1998) as human X-linked intellectual disability genes. Loss of a-PIX (Ramakers, Wolfer et al. 2012) or loss of both PAK1 and PAK3 (Huang, Zhou et al. 2011) in mice recapitulates this severe learning and memory phenotype. The direct evidence for a role for GIT1 in learning and memory is quite strong. Overexpressing GIT1 or interfering with GIT1 localization in primary hippocampal neurons alters dendritic spine density (Zhang, Webb et al. 2003). Memory defects have been reported in GIT1-deficient mice in aversive memory through fear conditioning (Schmalzigaug, Rodriguiz et al. 2009, Fass, Lewis et al. 2018), in associative memory using operant conditioning (Menon, Deane et al. 2010), in working memory through T-maze spontaneous alternation (Fass, Lewis et al. 2018) and in spatial and contextual memory in the Morris water maze and novel object recognition tests (Won, Mah et al. 2011, Martyn, Toth et al. 2018). In contrast, nothing has been reported concerning a direct role for GIT2 in learning and memory processes in neurons or in an animal model, although GIT2 has been presumed to be important in this function due to its similar biochemical functions compared to GIT1 (Premont, Claing et al. 2000, Zhou, Li et al. 2016, van Gastel, Boddaert et al. 2018)

One prominent report has linked a GIT1 gene polymorphism to attention deficit-hyperactivity disorder (ADHD) in a Japanese cohort, and genetrap mice lacking GIT1 were reported to model two critical aspects of ADHD: basal hyperactivity, and paradoxical calming by psychostimulants (Won, Mah et al. 2011). This linkage has been controversial, however. One group found no association of *GIT1* gene polymorphisms with human ADHD in a Brazilian population (Salatino-Oliveira, Genro et al. 2012), while another large study found no association in three large patient cohorts (Klein, van der Voet et al. 2015). Functionally, our group has shown that a distinct line of GIT1-deficient mice fails to demonstrate either hyperactivity or psychostimulant-induced locomotor suppression (Schmalzigaug, Rodriguiz et al. 2009, Martyn, Toth et al. 2018). Furthermore, a study using *Drosophila* also found no evidence for altered locomotor behavior in the absence of the single GIT gene, *dGIT* (Klein, van der Voet et al. 2015).

Here we have tested the hypothesis that GIT2-deficient mice might also exhibit an ADHD-like phenotype as well as learning and memory defects. Instead we show that GIT2-deficient mice are neither hyperactive nor display ADHD-like behaviors, but they also unexpectedly exhibit completely normal learning and memory function in several behavioral tests, as well as exhibit normal hippocampal long-term potentiation. This fundamental distinction between GIT2 and GIT1 in regulating learning and memory processes led us to examine GIT2 alternative splicing in the brain, since GIT2 is known to have extensive tissue-specific alternative splicing (Premont, Claing et al. 2000). We identify a predominant splice variant of GIT2 expressed in the brain that is lacking internal sequences that contain the coiled-coil region required for GIT dimerization. We find that this variant neither dimerizes nor forms tight oligomeric GIT/PIX complexes. The inability of the major brain form of GIT2 to function as part of oligomeric GIT/PIX scaffold complexes explains why loss of GIT2 in the brain does not affect synaptic PIX/PAK signalling required for learning and memory.

## Methods

### Plasmids

The pBK(Δ)-human GIT2-long/Flag plasmid has been described previously (Premont, Claing et al. 2000). This plasmid was used as template for site-directed mutagenesis to create GIT2(ΔBC)/Flag, GIT2(ΔE)/Flag and the double-deletion GIT2(ΔBCE)/Flag using the QuikChange mutagenesis kit (Stratagene). The ΔBC primers were 5’- ACTGCAAGCAAAACAAACCGGCAGAAGCTTCAAACACTCCAGAGTGAAAATTCG and 5’- GCAATTTTCACTCTGGAGTGTTTGAAGCTTCTGCCGGTTTGTTTTGCTTGCAGT to delete amino acids 415-464, and the ΔE primers were 5’-CCCTTCCCCGCGCACGCATCCAGGCTGGAG and 5’- CTCCAGCCTGGATGCGTGCGCGGCGAAGGG to delete amino acids 465-547, using the amino acid residue numbering of human GIT2-long.

### GIT2 variant PCR

To detect the internal alternative splicing of the GIT2 transcript, we prepared total RNA from dissected wildtype mouse brain regions using Qiazol (Qiagen) and prepared cDNA using the SuperScript III first-strand synthesis kit (Invitrogen). Primers spanning the A-B-C-D-E splicing region were 5’-GGTCAACCCTGAGTACTCCTC and 5’-AATCACTCTCCGGGGTGCTGT, and were used to amplify this region by PCR for 35 cycles. Resulting bands were isolated from an agarose gel and subjected to direct DNA sequencing using each of the amplification primers.

### Animals

The gene-trap *Git2* mice have been described previously (Schmalzigaug, Phee et al. 2007, Schmalzigaug, Rodriguiz et al. 2009), and were maintained on a mixed C57BL/6 x 1290la genetic background. All behavioral studies reported here used this gene-trap strain. A distinct second *Git2* knockout strain with a NEO insertion in exon 2 (Mazaki, Hashimoto et al. 2006) was obtained from Dr. Hisatake Sabe at 5 generations backcrossed to C57BL/6, and was further backcrossed to 12 generations C57BL/6J, but was used here only for brain GIT protein immunoprecipitation assays. Mice were housed 3-5/cage in a temperature- and humidity-controlled barrier facility on a 12h:12h light:dark cycle (lights on at 0700h). Chow diet and water were provided *ad libitum*. Behavioral assays used both male and female mice. All procedures were conducted with protocols approved by the Duke University Institutional Animal Care and Use Committee.

### Behavior

Testing for 24-hour locomotion, amphetamine-induced locomotion and learning and memory function using the Morris water maze and novel object recognition memory test were as described recently (Martyn, Toth et al. 2018). **Open field activity**. Motor activity in the open field was assessed in two separate experiments. In the first study, spontaneous locomotor activity was analyzed over 24 hours in a 42 x 42 x 30 cm open field (Omnitech Inc., Columbus, OH) illuminated at 340 lux. Activity was monitored in 30-min segments by 8 photobeams, spaced 2.5 cm apart, positioned 2.25 cm from the floor, and located around the perimeter of the open field (Martyn, Toth et al. 2018). Mice were placed into the apparatus at 1300 hr and removed 24 hr later. A second study evaluated locomotor responses to 0.5, 1, and 2 mg/kg amphetamine (Sigma-Aldrich, St. Louis, MO). Mice were placed into the open field for 60 min to assess baseline activity. They were removed, injected (i.p.) with AMPH, and returned immediately to the open field for 90 min. Locomotor activity was measured in 5-min segments and expressed as distance traveled in cm. **Fear conditioning**. Mice were placed into a MedAssociates fear conditioning apparatus (St. Albans, VT). After 2 min, a 30 sec 72 dB tone (CS) sounded that was terminated with a 2 sec 0.4 mA scrambled foot-shock (US); the mice remained in the conditioning apparatus for 30 sec and then were returned to their home-cage (Schmalzigaug, Rodriguiz et al. 2009, Porton, Rodriguiz et al. 2010). Twenty-four hr later the mice were tested in contextual fear by returning the mouse to the same chamber in which it had been conditioned in the absence of the CS and US for 5 min. The next day mice were tested for cued fear. They were placed into a novel chamber whose color, texture, shape, dimensions, and level of illumination were different from that of the conditioning chamber. After 2 min, the CS was presented for 3 min. All tests were videotaped and behaviors were scored by a trained observer blinded to the genotype of the mice using Noldus Observer software (Leesburg, VA). Freezing refers to the lack of all non-respiratory movement by the animal for >1 sec (Anagnostaras, Josselyn et al. 2000, Porton, Rodriguiz et al. 2010). **Novel object recognition memory**. Mice were trained by presentation of a pair of identical objects for 5 min and these objects constituted the “familiar” objects in the test. After 20 min mice were tested for short-term (STM) and were tested for long-term memory (LTM) 24 hr after training. In each case, a single familiar object was paired with a novel object. All behaviors were filmed and were scored subsequently with Noldus Ethovision by observers who were blind to the genotypes and sex of the animals. Preference scores were calculated by subtracting the total time spent with the familiar object from time spent with the novel object, and dividing this difference by the total amount of time spent with both objects. Positive scores indicated preferences for the novel object, negative scores denoted preferences for the familiar object, and scores approaching “zero” signified a preference for neither object. **Spatial learning and memory in the Morris water maze**. All training and testing were conducted in a 120 cm diameter pool, maintained at 24°C, and under ~125 lux illumination. The pool was divided into northeast (NE), northwest (NW), southeast (SE) and southwest (SW) quadrants. Prior to testing, mice were handled, acclimated to standing in water, and trained to sit on and swim around the hidden platform. Testing was divided into 2 phases: acquisition (days 1-8) with the hidden platform in the NE quadrant and reversal (days 9-16) with the hidden platform in the SW quadrant. Mice received 4 trials a day in pairs that were separated by 60 min. Release points were randomized across trials and days. Every other day, a single probe trial where the platform was removed from the maze was given 1 hr after the 4 test-trials. The same cohort of mice was used for visible platform testing with 4 trials a day over 5 consecutive days. For this test, mice were released from the point opposite the platform and given 60 sec to swim to the visible platform. The platform location was changed on each trial to a new, randomized location. All trials ended when the animal reached the platform or after 60 sec had elapsed. Performance was filmed by high-resolution camera suspended 180 cm above the center of the pool and scored by blinded observers using Ethovision XT 7 (Noldus). Tracking profiles were generated and were used to measure swim time and swim velocity.

### Spine density

For determination of spine morphology and density, eleven day-old mice were decapitated following an overdose with Nembutal. Slice cultures were prepared as described previously (Simons, Escobedo et al. 2009). Briefly, the whole brain was removed under sterile conditions and immersed in ice cold MEM (GIBCO Technologies) supplemented with 25 mM Hepes, 10 mM Tris-base, 10 mM glucose, and 3 mM MgCl_2_. Slices containing hippocampus were then cut at 200 μm on a vibrating tissue slicer (Leica) and placed into the center of a membrane in a transwell plate (Costar). Culture media was prepared as a 2:1 mixture of Basal Medium Eagle (Sigma) and Earle’s Balanced Salts Solution (Sigma), respectively, and supplemented with 20 mM NaCl, 5 mM NaHCO_3_, 0.2 mM CaCl_2_, 1.7 mM MgSO_4_, 48 mM glucose, 26.7 mM Hepes, 5% horse serum (GIBCO), 10 ml/liter penicillin-streptomycin (GIBCO), 1.32 mg/liter insulin (Sigma), and the pH adjusted to 7.2. The slices were incubated in 5% CO_2_ at 34°C and half the media replaced daily. On DIV 6-7, the organotypic slices were infected with a recombinant Sindbis virus for expression of EGFP. Infected slices with sufficient EGFP expression were fixed with 4% paraformaldehyde and imaged using an Axioskop 2FS microscope (Carl Zeiss, Inc., Thornwood, NY) coupled to a Zeiss LSM 510 NLO META system and a Ti:sapphire Chameleon two-photon laser system. Images of dendrites in the CA1 region of the hippocampus were acquired with a two-photon laser tuned to 900 nm (Coherent, Inc., Auburn, CA) for high resolution, and an Argon laser at 488nm (for eGFP excitation) for larger views of the slice. Individual Z-stack images containing 35-40 micron lengths of dendritic segments from 17 WT and 18 KO slices from 5 mice for each genotype were analyzed using the Zeiss software for spine density and morphology according to the method outlined by (Chapleau, Carlo et al. 2008). Briefly, a protrusion was considered to be a spine if it extended less than or equal to 3μm from the parent dendrite. Each spine was counted only once by following its projection course through the stack of z sections. Spine density was calculated by quantifying the number of spines per dendritic segment length. For each spine, morphology was determined by measuring the diameter of the neck of each spine (N), its length (L), and the diameter of each head (H). Spines were classified into 3 types: stubby, mushroom and thin, based on the ratios of L/N and H/N. Stubby spines have L, N, and H dimensions that are all similar to each other (L~N~H). Mushroom spines have an H/N ratio greater than one, with H>N. Thin spines have L>N. WT and GIT2-KO spine density were compared using the Mann-Whitney U (Rank sum test).

### Electrophysiology

Slices of hippocampus were prepared as described (Bastrikova, Gardner et al. 2008). Animals were deeply anaesthetized with Nembutal and decapitated prior to brain removal. Slices of hippocampus were cut using a vibrating tissue slicer (Leica) in sucrose-substituted artificial cerebral spinal fluid (ACSF) in mM: 240 sucrose, 2.0 KCl, 1 MgCl_2_, 2 MgSO_4_, 1 CaCl_2_, 1.25 NaH_2_PO_4_, 26 NaHCO_3_, and 10 glucose, which was bubbled with 95% O2/5% CO_2_. Slices were transferred directly to an interface-type recording chamber where they were allowed to incubate for at least one hour before recording. Slices were continuously bathed at 34° C with standard ACSF (in mM): 124 NaCl, 2.5 KCl, 2 MgCl_2_, 2 CaCl_2_, 1.25 NaH_2_PO_4_, 26 NaHCO_3_, and 17 D-glucose. Synaptic responses in the CA1 region were evoked with a bipolar stimulating electrode placed in the stratum radiatum of CA1 and recorded with an ACSF-filled glass pipette, also placed in the stratum radiatum. Long-term potentiation (LTP) of the field excitatory post-synaptic potential (fEPSP) was induced with three episodes of theta burst stimulation (TBS; ten 100 Hz bursts of 4 pulses delivered at 5 Hz, 30 seconds apart). Partial depotentiation was induced with 900 pulses delivered over 7.5 minutes at 2 Hz (low-frequency stimulation; LFS). Data are expressed as a percentage of the average baseline response collected in the fifteen minutes prior to the TBS, and are presented as an mean from 15 slices from 9 mice +/− SEM. Paired-pulse frequency (PPF) was measured by delivering two electrical pulses in short succession and was expressed as a ratio of the size of the second synaptic response to the size of the first synaptic response.

### Cell culture, transfection, immunoprecipitation and western blotting

COS7 cells were maintained at 37°C under 5% CO_2_ in DMEM media (Life Technologies) supplemented with 10% fetal bovine serum (Atlanta Biologicals), 100 U/mL penicillin and 100 μg/mL streptomycin (Life Technologies). Cells were transfected with plasmid DNA using Polyfect (Qiagen), according to the manufacturer’s instructions. Two days after transfection, cells were scraped into lysis buffer [50mM Tris-HCl pH 8.0, 150mM NaCl, 0.5% (v/v) Triton X-100, 0.5% (v/v) NP-40, 0.5% (w/v) deoxycholate, 0.1% (w/v) SDS] supplemented with protease inhibitor cocktail (Sigma), rotated for 1 hour at 4°C, and pelleted at 21,000xg for 20 minutes at 4°C. Solubilized lysate was immunoprecipitated using M2-Flag-agarose conjugate (Sigma). Protein samples were separated using 10% polyacrylamide gels (BioRad), and transferred to nitrocellulose for immunoblotting. Signals were detected on X-ray film using ECL chemiluminescence reagent (GE). Anti-GIT1 H-170 antiserum was from Santa Cruz, GIT1 and PKL monoclonal antibodies were from Becton-Dickinson, p50 β-PIX antiserum was the kind gift of Dr. Rick Cerione (Cornell University), and M2 Flag-peroxidase conjugate and M2 Flag-agarose beads were from Sigma. Secondary anti-rabbit-HRP and anti-mouse-HRP were from GE, and TrueBlot anti-mouse-HRP was from Rockland.

### Immunoprecipitation of native GIT2 and GIT1 from brain lysates

An entire mouse brain was homogenized in 10 volumes of lysis buffer with 20 strokes in a Dounce homogenizer, rotated for 1 hour at 4°C, and pelleted at 21,000xg for 20 minutes at 4°C. Soluble lysate (1 ml) was immunoprecipitated overnight using 2 μg of PKL monoclonal antibody, which was raised against chicken GIT2 but recognizes both GIT2 and GIT1 from mammals. Immune complexes were captured with Protein G/Protein A plus-agarose (Calbiochem) and subjected to Western blotting using PKL antibody (25 ng/ml) and anti-mouse TrueBlot-HRP secondary. Brains used were from WT and genetrap GIT2-KO mice, but also GIT1-KO (Schmalzigaug, Rodriguiz et al. 2009) and a distinct line of GIT2-KO mice made using a traditional NEO replacement strategy (Mazaki, Hashimoto et al. 2006).

### Statistical analysis

Data were analyzed by one-way or repeated measures ANOVA test for comparison between genotypes, treatments, or doses (GraphPad Prism 6 software). Individual genotypes, treatments, or doses were compared using a post-hoc test as indicated in the figure legends whenever ANOVA showed significance to either genotype or genotype x time interaction. A probability value of p<0.05 was considered as statistically significant. All data are presented as mean ± SEM.

## Results

### GIT2-KO mice do not exhibit ADHD-like behavior

A prominent report suggested that mice lacking GIT1 exhibited behavioral abnormalities consistent with an Attention Deficit-Hyperactivity Disorder (ADHD)-like phenotype (Won, Mah et al. 2011). These included basal hyperactivity and psychostimulant-induced locomotor suppression rather than activation, as well as learning and memory deficits. GIT2 has not been examined previously for ADHD-like behavior, although we previously reported that GIT2-KO mice displayed sex-dependent differences in locomotor activity in the first 5 minutes in a novel chamber, consistent with elevated anxiety in female GIT2-deficient mice (Schmalzigaug, Rodriguiz et al. 2009). We therefore examined spontaneous activity of GIT2-deficient mice in more detail by recording locomotor activity over a complete 24-hour diurnal cycle (Fig 1). Following the initial habituation period, GIT2-deficient mice exhibited low activity in the light and higher activity in the dark, but overall levels of activity in either light or dark did not differ significantly by genotype, comparing knockout to wildtype littermates.

**Figure 1.**
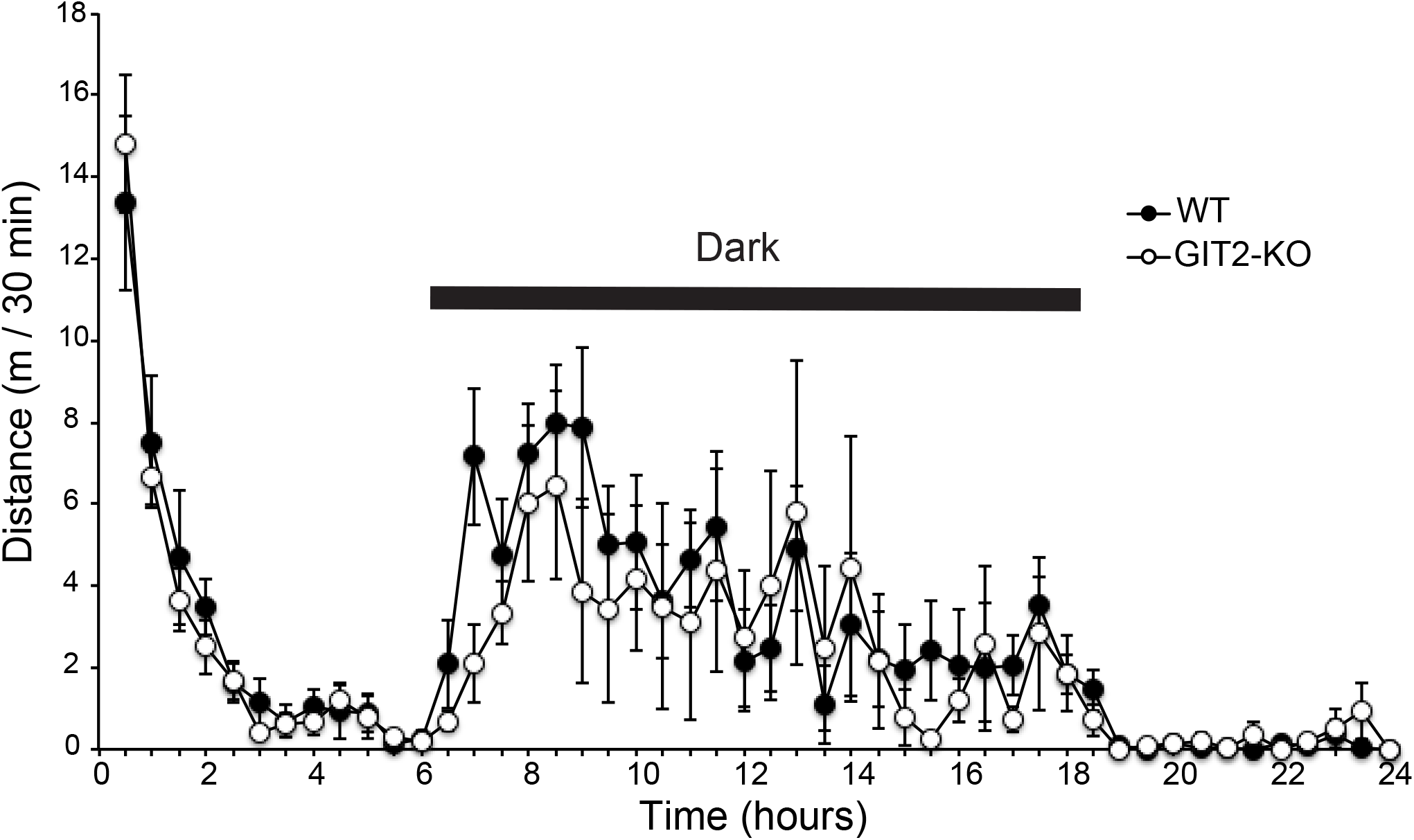
Spontaneous locomotor activity in GIT2 genetrap mice. A) GIT2 WT (n=5, black circle) and GIT2 KO (n=5, open circle) mice were placed in the open field for 24 hours under the normal 12:12hr light:dark cycle. Distance traveled (m) in each 30 min period is shown. Statistical analysis using repeated measures ANOVA showed no significant difference between genotypes for activity measured in the dark or in the light over the entire test period.

The paradoxical motor calming effect of psychostimulants in ADHD patients is utilized therapeutically, and was reported to occur in mice with a genetrap inactivation of the *Git1* gene following administration of amphetamine or methylphenidate (Won, Mah et al. 2011). We tested the acute locomotor responses to amphetamine at three doses in GIT2-deficient mice (Fig 2). In no case did amphetamine provoke locomotor suppression. At low amphetamine (0.5 mg/kg), GIT2-deficient mice responded with significant locomotor activation while WT mice did not respond significantly to that dose (Fig 2A,B), while at 1 or 2 mg/kg, both genotypes responded equivalently over the 2h test period (Fig 2B,C and 2D,E). These data suggest that mice lacking GIT2 have increased sensitivity to amphetamine compared to WT, but show that the drug provokes typical locomotor stimulatory effects in these mice rather than locomotor suppression such as is observed in ADHD patients.

**Figure 2.**
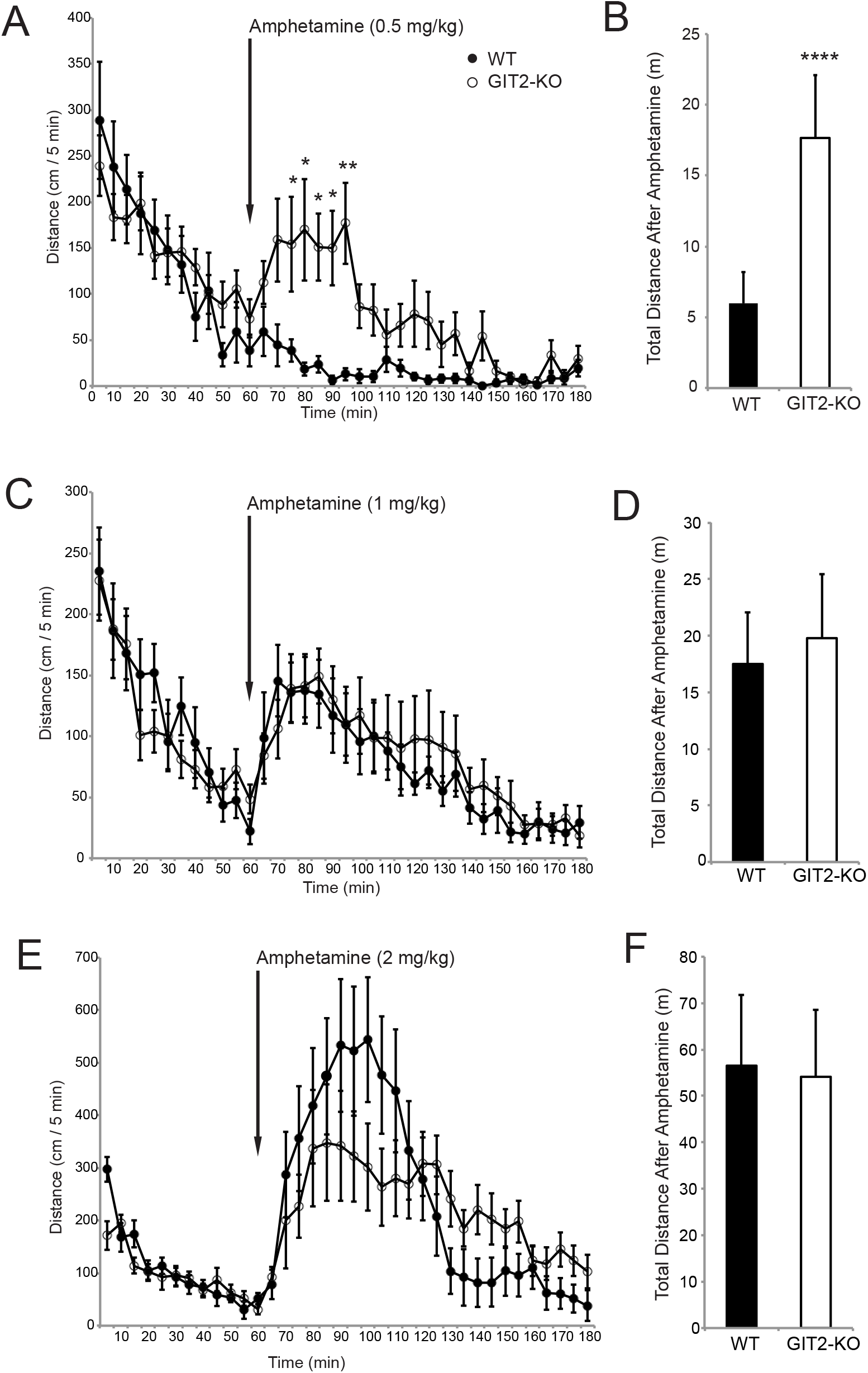
Amphetamine-induced locomotor activity in GIT2 genetrap mice. GIT2 WT (black circle) and GIT2 KO (open circle) mice were habituated to the locomotor chamber for 60 min prior to drug injection. A) GIT2 WT (n=10) and KO (n=9) mice were injected with 0.5mg/kg amphetamine, and locomotor activity is shown in 5 min windows. *p*<0.001 in a two-way repeated measures ANOVA between genotypes over time; * *p*<0.05, ** *p*<0.01 within time using a Holm-Sidak post-hoc test. B) Total locomotor activity summed for the 2h following 0.5mg/kg amphetamine injection. **** *p*<0.001 using t-test. GIT2 WT (n=9) and KO (n=11) mice were injected with 1mg/kg amphetamine, and are shown as locomotor activity in 5 min windows (C) or summed over 2h after drug (D). No significant differences. GIT2 WT (n=10) and KO (n=10) mice were injected with 2mg/kg amphetamine, and are shown as locomotor activity in 5 min windows (E) or summed over 2h after drug (E). No significant differences.

### GIT2-KO are not deficient in fear conditioning

A distinct aspect of ADHD, reduced attention and focus, was reported in GIT1-deficient mice as reduced performance in learning and memory tests (Won, Mah et al. 2011). Indeed, several reports have shown multiple learning and memory deficits in global GIT1-knockout mice (Schmalzigaug, Rodriguiz et al. 2009, Menon, Deane et al. 2010, Won, Mah et al. 2011, Martyn, Toth et al. 2018) and in neuron-specific GIT1-KO mice (Fass, Lewis et al. 2018). Mice lacking GIT2 have not been tested previously for cognitive function, so we assessed their learning and memory behavior using specific tests.

Aversive memory was tested using a classical conditioning paradigm, auditory fear conditioning (Rodriguiz and Wetsel 2006). Mice were introduced to a novel chamber, and after 2 minutes of acclimation, a tone was sounded for 30 seconds, terminating with 2 seconds of scrambled footshock (Fig 3A). Fear was assessed as behavioral freezing, and GIT2-KO showed significantly elevated freezing behavior immediately after the tone/shock. On the day after conditioning, mice were reintroduced to the same chamber, with no tone and no shock, and both GIT2-KO and wildtype mice exhibited a comparably high degree of freezing behavior indicative of normal conditioned memory of the prior conditioning context (Fig 3B). On the following day, mice were introduced into a novel chamber, and after 3 minutes, the tone was sounded for 2 minutes but no shock was presented. GIT2-KO mice exhibited a notable degree of freezing prior to the presentation of the tone, compared to the WT mice, but both genotypes responded robustly when the tone was presented (Fig 3C). Thus, like WT mice, GIT2-KO mice remember the aversive conditioning to both the shock context (the original chamber) and to the conditioned tone in a distinct context.

**Figure 3.**
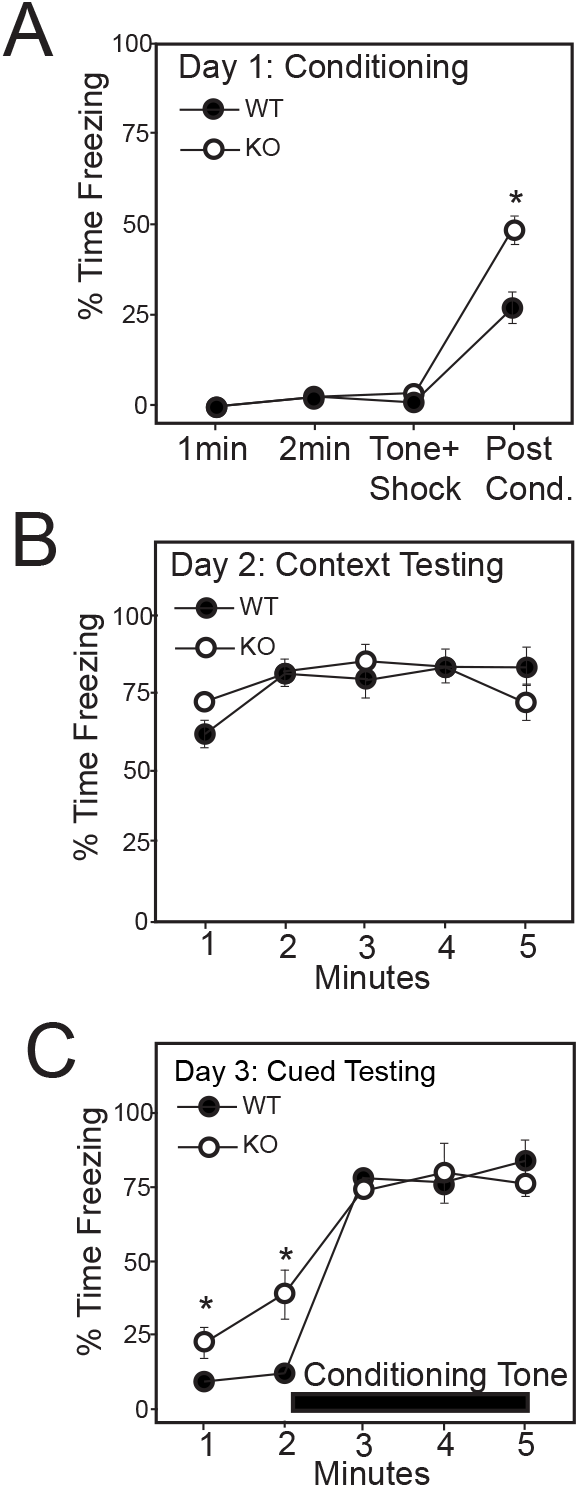
Aversive learning by auditory fear conditioning in GIT2-KO mice. GIT2 WT (n=9, black circle) and GIT2 KO (n=9, open circle) were subjected to auditory fear conditioning and tested for freezing behavior on day 1 (A), tested context-dependent freezing behavior on day 2 (B), and tested for cue (tone)-dependent freezing on day 3 by presenting the conditioning tone from min 2-5 (black bar) (C). \**p*<0.05, WT versus KO mice using repeated measures ANOVA and post-hoc Sidak Multiple Comparison test.

### Female GIT2-KO mice are selectively deficient in episodic memory

Short- and long-term object recognition memory were measured using the novel object test (Rodriguiz and Wetsel 2006). Mice were acclimated to a test arena containing two identical objects, and then were tested for object memory by replacing one initial object with a distinct novel object. Since rodents prefer to examine a new object rather than a previously encountered one, the number of object contacts and time spent interacting with each object was measured to calculate a preference ratio (Figure 4). At training, neither genotype demonstrated any preference for one identical object over the other, but mice lacking GIT2 exhibited a significant sex difference in subsequent testing and were analyzed separately. Male GIT2-KO mice were indistinguishable from their male WT controls, and preferred to interact with the novel object when tested 20 min after training (STM) and when tested 24 hr later (LTM) (Figure 4A). In contrast, female GIT2-KO mice selected the familiar over the novel object in both the STM and the LTM tests, whereas the WT females strongly preferred the novel object (Figure 4B), suggestive of neophobia in GIT2-KO females. As a control, the numbers of object contacts were compared for both male and female mice, and neither differed from wildtype (Figure 4A,B). This neophobia in GIT2-KO females is consistent with the elevated anxiety phenotype of GIT2-KO mice, particularly females (Schmalzigaug, Rodriguiz et al. 2009). Nonetheless, the female GIT2-KO clearly distinguished between familiar and novel objects, indicative of effective learning of the familiar object. Collectively, these findings show that mice lacking GIT2 are able to learn to differentiate between novel and previously encountered objects.

**Figure 4.**
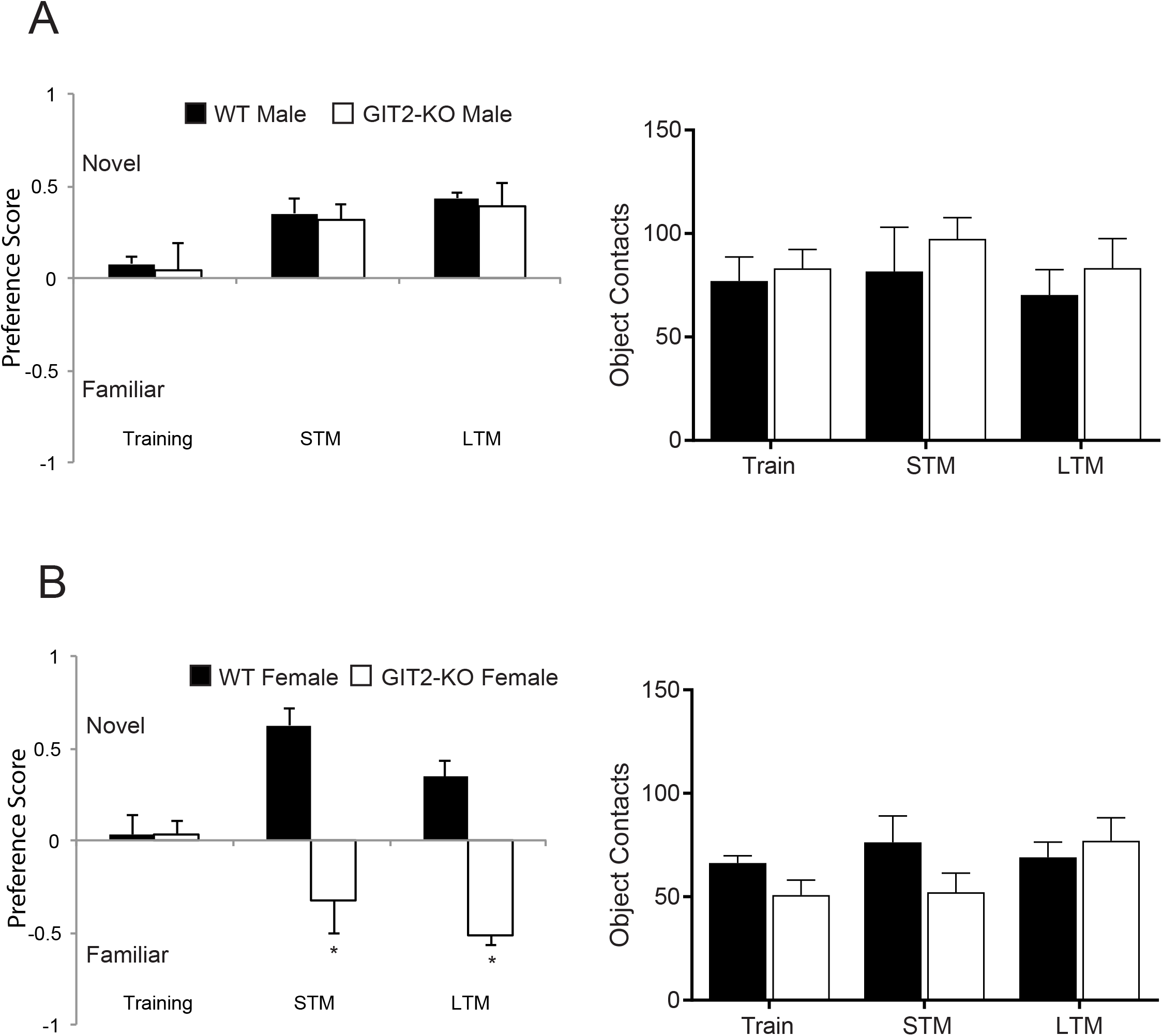
Novel object recognition memory in GIT2 KO mice. A) Male GIT2 WT (n=6, black bars) and GIT2 KO (n=7, open bars) mice were trained using two identical objects, and then tested for short-term memory (STM) and long-term memory (LTM) using one novel object in place of one previously presented object. Male GIT2 WT and KO mice appeared indistinguishable during training and in showing preference for exploring the novel object at both test times, and had similar numbers of object contacts in all test periods. No significant differences between genotypes were detected at each test period using one-way ANOVA and Tukey’s Multiple Comparison test. B) Female GIT2 WT (n=6, black bars) and GIT2 KO (n=6, open bars) mice appeared indistinguishable during training but differed in both short-term and long-term memory test. Wildtype female mice preferred the novel object, while female GIT2 KO mice preferred the familiar object; nevertheless, both genotypes had similar numbers of object contacts. * *p*<0.05 for genotype using one-way ANOVA and Holm-Sidak post-hoc test.

### GIT2 KO mice show normal spatial learning and memory

Spatial learning was assessed using the Morris water maze, where mice are trying to escape the water and have the learn the location of the hidden platform. (Rodriguiz and Wetsel 2006). GIT2-KO and WT mice reduced their total distance to locate the hidden platform on successive days, and did not differ between genotypes, indicative of normal spatial learning behavior (Fig 5A). After the acquisition-learning phase, the platform was moved to a new location, and reversal learning was assessed. GIT2-KO and WT mice rapidly learned the new location, as assessed by swim distance, and did not differ by genotype. During acquisition and reversal learning, probe trials with the platform absent were conducted to examine the evolution of search strategy over time, and during both acquisition and reversal training, both WT and GIT2-KO mice increasingly spent more time in the quadrant that had contained the platform and less time in other quadrants, and did not differ by genotype (Fig 5B,C). GIT2-KO swam significantly more slowly than WT mice in both test phases (Fig 5D), but not slowly enough to affect interpretation of the results as a learning paradigm. As a control, the visible platform variant of the water maze test was performed, and both GIT2-KO and WT controls rapidly found the platform (Fig 5E), but the GIT2-KO continued to demonstrate reduced swimming speed (Fig 5F).

**Figure 5.**
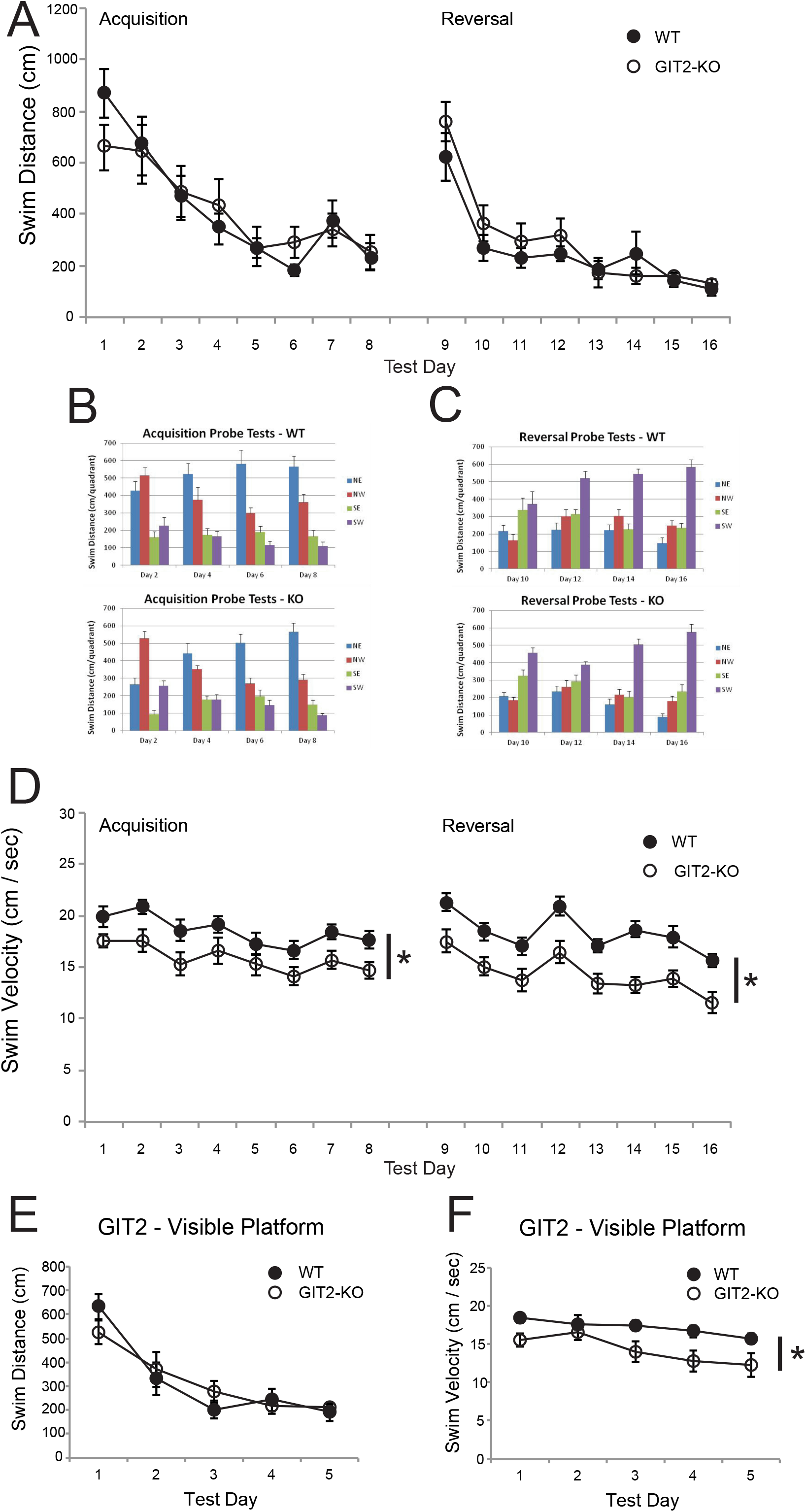
Spatial learning and memory in GIT2 KO mice in the Morris water maze. A) Swim distance to hidden platform during acquisition learning (days 1-8) and reversal learning (days 9-16), GIT2 WT (n=10, black circles) and GIT2 KO (n=10, open circles). No significant difference between genotypes. B) Time spent swimming in each quadrant during a probe trials where the platform was absent, during alternate days following acquisition training (days 1-8). Mice exhibit increased time in the quadrant that had contained the platform (NE) and decreasing time in other quadrants across days. No significant differences between genotypes. C) Time spent swimming in each quadrant during a probe trials where the platform was absent, during alternate days following reversal training (days 9-16). Mice exhibit increased time in the quadrant that had contained the platform (SW) and decreasing time in other quadrants across days. No significant differences between genotypes. D) Swim velocity during initial learning and reversal learning was significantly higher in GIT2-deficient mice over test days by repeated measures ANOVA, * *p*=0.006 for acquisition and * *p*=0.003 for reversal learning. E) Visible platform test, swim distance during initial learning (days 1-5), WT (n=8, black circles) and KO (n=9, open circles). No significant difference between genotypes. F) Swim velocity during acquisition learning in the visible platform test over test days by repeated measures ANOVA, * *p*=0.03.

### GIT2-KO reduces hippocampal dendritic spine density without affecting LTP

Altered learning behavior often is associated with reduced dendritic spine density and with spine immaturity, in human intellectual disability patients and in mouse models exhibiting poor learning (Levenga and Willemsen 2012, Ba, van der Raadt et al. 2013). The GIT partner a-PIX and the GIT/PIX partner PAK3 are both known human X-linked intellectual disability genes, and are thought to affect the same pathway (Allen, Gleeson et al. 1998, Kutsche, Yntema et al. 2000, Ba, van der Raadt et al. 2013). Mice lacking a-PIX (Ramakers, Wolfer et al. 2012) or PAK1 plus PAK3 (Huang, Zhou et al. 2011), exhibit profound learning deficits together with altered dendritic spine density, as do GIT1-deficient mice (Menon, Deane et al. 2010, Martyn, Toth et al. 2018). We therefore examined the density of hippocampal dendritic spines in the absence of GIT2. We used 2-photon microscopy to image cultured brain slices from post-natal pups that were infected with EGFP in order to assess both spine density and spine morphology. Hippocampal CA1 neurons in brain slices from GIT2-knockout mice demonstrated a reduced density of dendritic spines (Fig 6A,B). However, analysis of spine morphology revealed that the distribution of thin, mushroom and knobby spines was not altered by the absence of GIT2 (Fig 6C). Thus GIT2 does appear to regulate dendritic spines (perhaps by affecting the probability of formation or spine stability), but does not appear to affect maturation *per se* as assessed by the distribution of morphological types.

**Figure 6.**
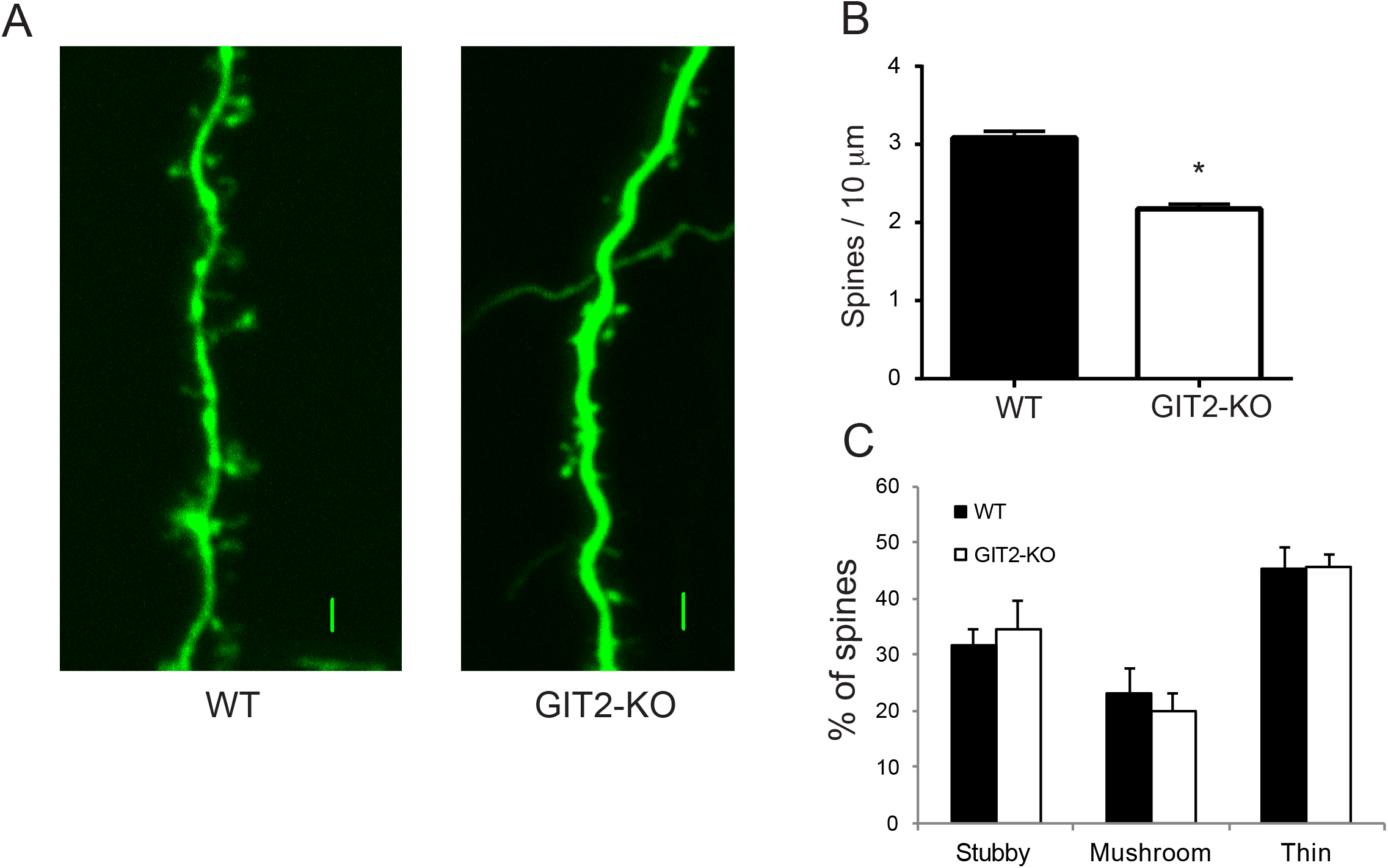
Hippocampal CA1 synaptic spines are reduced in GIT2-deficient mice. A) Representative fluorescent micrographs of GFP-infected CA1 neurons showing spines. Scale bars are 2 μm. B) Spine density was calculated by dividing the number of spines by the length (μm) of the dendritic segment for each image counted, using 76 and 85 images from 5 WT mice and 5 GIT2-KO mice, respectively. * *p*<0.005 using t-test. C) When counted, each individual spine was categorized as stubby, mushroom or thin based on length and width, and spine morphology distribution is presented as % of spines with each morphology type. Spine morphology distribution was not significantly altered by loss of GIT2.

To directly assess synaptic plasticity, brain slices from GIT2-knockout mice were subjected to electrophysiological recording of hippocampal CA1 neurons to measure long-term potentiation of glutamate-induced excitatory synaptic currents. Consistent with the normal performance of GIT2 knockout mice in learning and memory tests, but surprisingly in light of reduced GIT2-KO spine number, hippocampal CA1 neurons from GIT2-deficient mice exhibited normal LTP (Fig 7).

**Figure 7.**
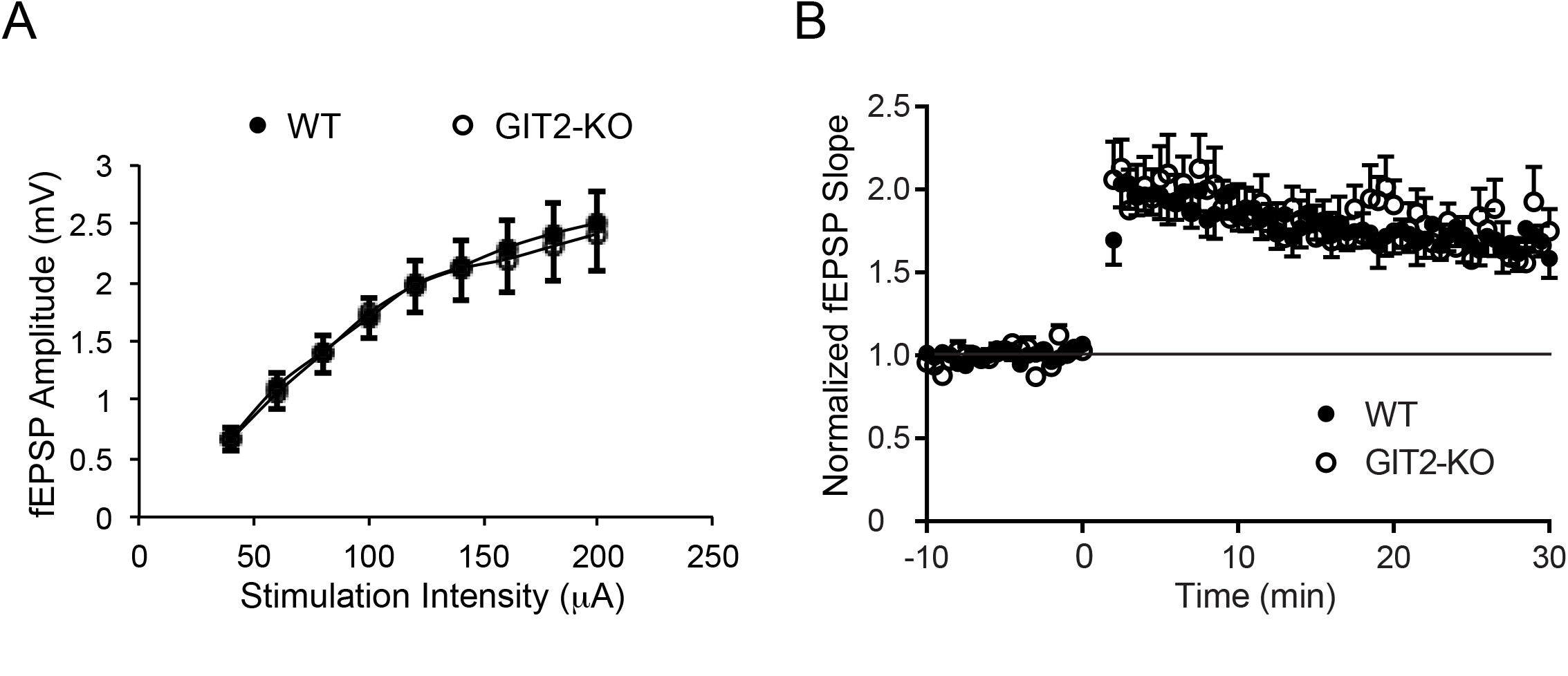
Hippocampal CA1 long-term potentiation is not altered by loss of GIT2. Hippocampal slices from WT and GIT2-KO mice (15 slices from 9 mice of each genotype) were stimulated at time 0 with 3 episodes of theta burst stimulation (TBS) to induce LTP. Data are expressed as percent of average baseline EPSP slope response, and are presented as mean ± SEM. No significant differences were observed.

### GIT2-KO differs from GIT1-KO due to the inability of brain GIT2 to form GIT/PIX complexes

Overall, the behavioral phenotypes of GIT1-deficient versus GIT2-deficient mice appear quite distinct. Loss of GIT1 leads to poor learning and memory behavior and is associated with reduced synaptic structural plasticity, whereas loss of GIT2 leads to elevated anxiety but has no significant effect on learning and memory and is associated with normal synaptic plasticity. This is unexpected, since GIT1 and GIT2 are widely expressed throughout the brain (Schmalzigaug, Phee et al. 2007), are capable of heterodimerizing within GIT/PIX complexes in cells that co-express the two isoforms (Premont, Perry et al. 2004), and share Arf GAP function and multiple protein partners (Premont, Claing et al. 2000, Zhou, Li et al. 2016). Loss of GIT1 in the brain leads to a substantial reduction in PIX levels (Won, Mah et al. 2011), and a recent report demonstrated a similar loss of PIX in immune tissues from GIT2-deficient mice, and showed that this was a result of destabilization of GIT-free PIX rather than altered gene transcription (Hao, He et al. 2015). However, immunoblotting for β-PIX in hippocampal lysates from GIT2-KO and WT mice revealed that PIX levels are *not* reduced in the absence of GIT2, while we confirm that PIX proteins are reduced dramatically in the absence of GIT1 (Fig 8). There also was no apparent compensatory up-regulation of GIT1 expression in GIT2-KO mice, which might have explained why PIX levels remain high. This suggests that brain GIT2 must somehow act quite differently from brain GIT1 or immune cell GIT2 with regard to complexing with and stabilizing PIX proteins.

**Figure 8.**
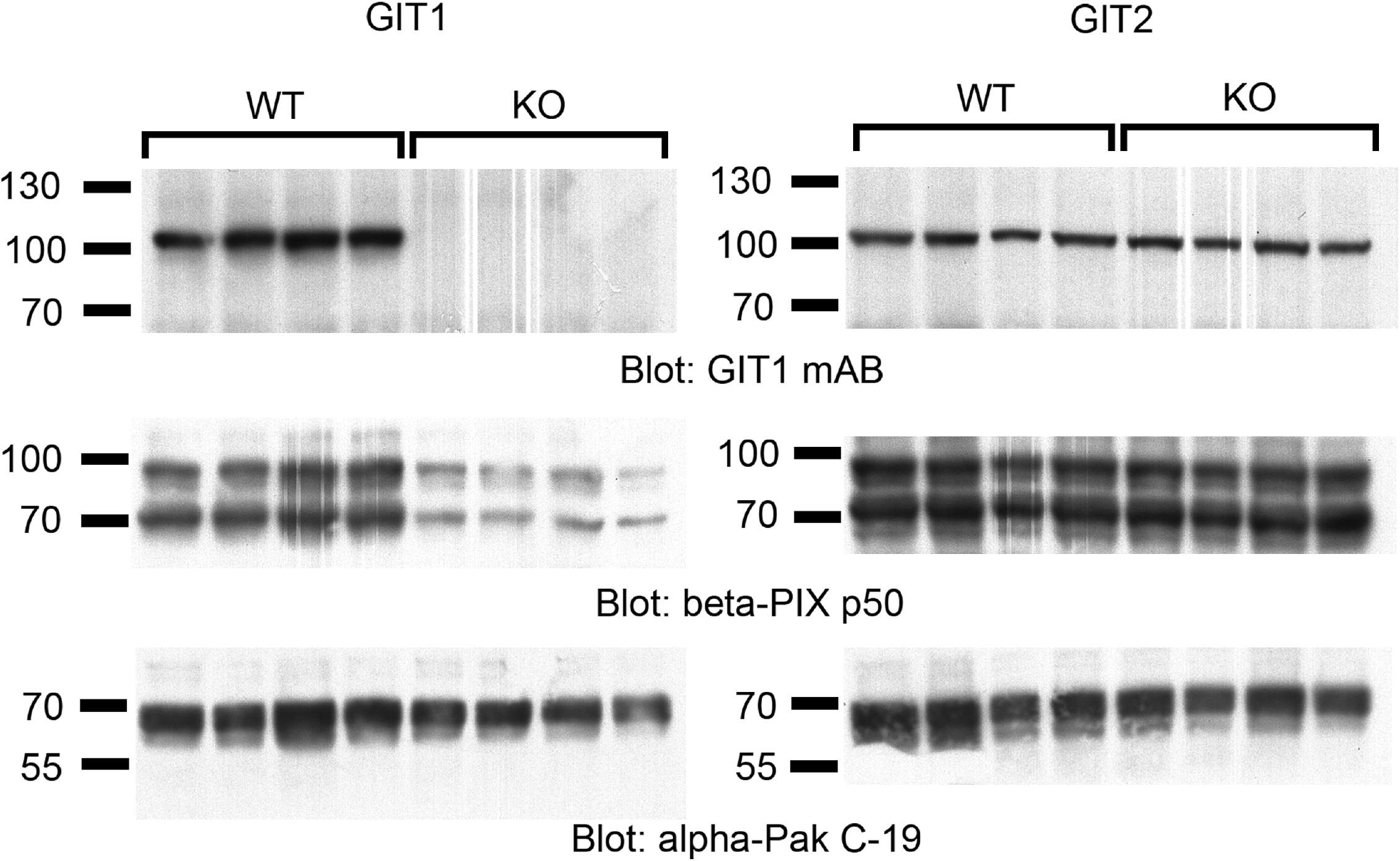
β-PIX, GIT1 and PAK levels are unaltered in GIT2 KO mouse hippocampus, but β-PIX levels are reduced in GIT1 KO hippocampus. Western blot analysis was performed on hippocampal protein lysates from individual GIT2 WT (n=4), GIT2 KO (n=4), GIT1 WT (n=4) and GIT1 KO (n=4) mice using GIT1 H-170, β-PIX p50, α-PAK (PAK1) and β-PAK (PAK3) antibodies, respectively. Blots shown are representative of two experiments.

To better understand the mechanistic basis for these fundamental differences between GIT2 and GIT1 function, we examined GIT2 in the brain in more detail. Specifically, from the first description of GIT2, it has been known that the *Git2* transcript undergoes extensive, tissue-specific alternative mRNA splicing of five contiguous internal sequences (encoded by 4 in-frame exons) leading to potentially over 30 variants (Premont, Claing et al. 2000). Comparative characterization of the longest and shortest GIT2 forms demonstrated that many properties are common to both variants, including Arf GAP activity and formation of GIT/PIX complexes (Premont, Claing et al. 2000). However, other GIT2 isoforms have never been compared directly. To identify the GIT2 splice variants expressed in the brain, total RNA was isolated from mouse hippocampus and amygdala and used in reverse transcription PCR with a primer pair spanning the alternatively spliced region, with an expected product of 820bp for the longest form of GIT2. A single product band of 570bp was obtained in both cases, and these were excised and subjected to DNA sequencing using the two amplification primers. Comparison of the resulting sequences to full-length mouse GIT2 revealed a loss of two blocks of sequences, corresponding to regions B and C together (coding exon 14) and region E (coding exon 16) (Fig 9A). This variant is described from mouse and several other species (including human) in the GenBank database as GIT2 isoform 3. Aligning this GIT2(ΔBCE) variant with the previously characterized GIT2-long and GIT2-short, it is apparent that the BC region contains most of the coiled-coil sequence that mediates GIT protein dimerization (Kim, Ko et al. 2003, Paris, Longhi et al. 2003, Premont, Perry et al. 2004, Schlenker and Rittinger 2009). The effect of the loss of this BC region has not been reported.

**Figure 9.**
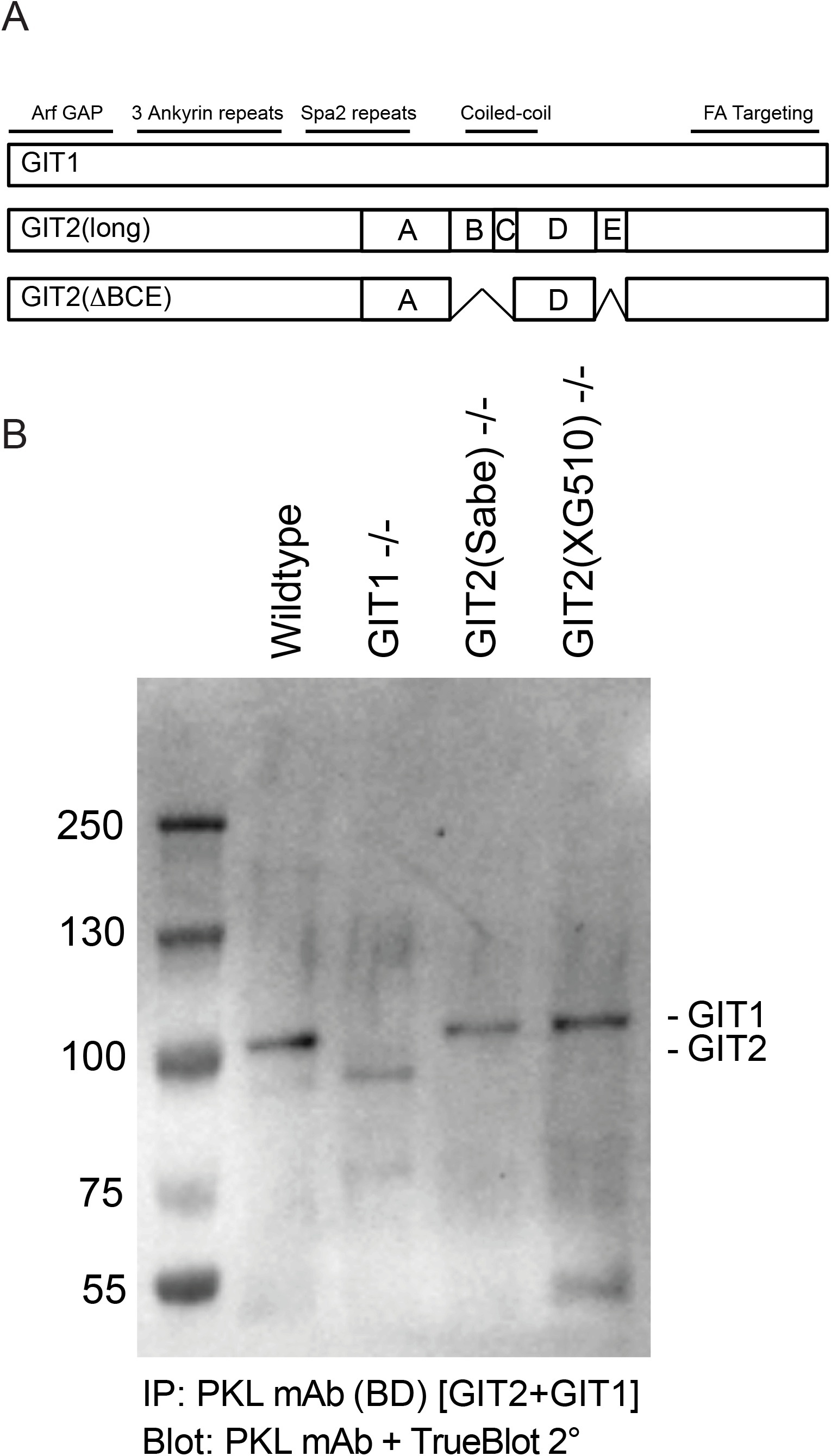
Brain GIT2 is predominantly a splice variant lacking two internal regions. A) Schematic diagram of GIT2 variants due to mRNA splicing, indicating the BC and E regions found to be missing in GIT2 mRNA amplified from mouse hippocampus and amygdala. The locations of functional domains are indicated. B) Brain GIT2 unambiguously identified by immunoprecipitation of GIT1+GIT2 using a non-selective antibody from whole brain lysates from WT, GIT1 KO and GIT2 KO mice. Brain GIT1 is a single 95 kDa band that is absent in GIT1 KO, while the predominant brain GIT2 band is not at 95 kDa as expected for full length GIT2 but is present at a smaller size (~87 kDa), and is absent in two distinct GIT2-KO strains. GIT2(XG510) is the GIT2 genetrap-KO line characterized behaviorally here, while GIT2(Sabe) is the traditional NEO replacement knockout line described by the Sabe lab (Mazaki, Hashimoto et al. 2006). Blots shown are representative of two independent experiments.

The longest form of GIT2 is co-linear with GIT1, and recombinant GIT2-long migrates at the same apparent size as GIT1 on SDS-PAGE (Premont, Claing et al. 2000). In our hands, the quality of anti-GIT2 antisera available commercially, or of several sera we have raised ourselves, is inadequate to cleanly detect native brain GIT2 without also cross-reacting with GIT1 or detecting contaminant bands that do not disappear in knockout samples. Thus, few groups have reported clear detection of native GIT2 variants without simultaneous detection of GIT1, particularly with unambiguous knockout or antigen-block controls (Schmalzigaug, Rodriguiz et al. 2009, Totaro, Tavano et al. 2012). To circumvent this problem, we utilized the PKL (chicken GIT2) monoclonal antibody, which strongly binds to both GIT2 and GIT1, to immunoprecipitate the native GIT proteins from wildtype or GIT knockout brain for subsequent Western blotting (Fig 9B). In wildtype brain, a strong 95 kDa band and a weaker 85 kDa band are seen. PKL antibody immunoprecipitates from GIT1-KO brain lack the 95 kDa band, while PKL antibody immunoprecipitates from brains of two distinct GIT2-KO strains (our own genetrap line and the traditional NEO replacement line from the Sabe lab (Mazaki, Hashimoto et al. 2006)) lack the lower p85 band. This unambiguously identifies the upper p95 band as GIT1 and the lower p85 band as GIT2, and demonstrates that the predominant form of GIT2 in the mouse brain is a molecular form that is substantially shorter than GIT1 or GIT2-long, consistent with our RNA amplification data identifying GIT2(ΔBCE). There appears to be very little GIT2-long protein (that is, migrating at the same apparent size as GIT1) in the mouse brain.

We created the human GIT2(ΔBCE) expression construct, and used this to examine the ability of this brain form of GIT2 to dimerize (Fig 10A). GIT2(ΔBCE) was well expressed, and as expected exhibited notably faster migration in SDS-PAGE compared to GIT2-long. In contrast to the clear dimerization ability of GIT2-long, GIT2(ΔBCE) lacks the ability to dimerize with and thus strongly co-immunoprecipitate with native GIT1.

**Figure 10.**
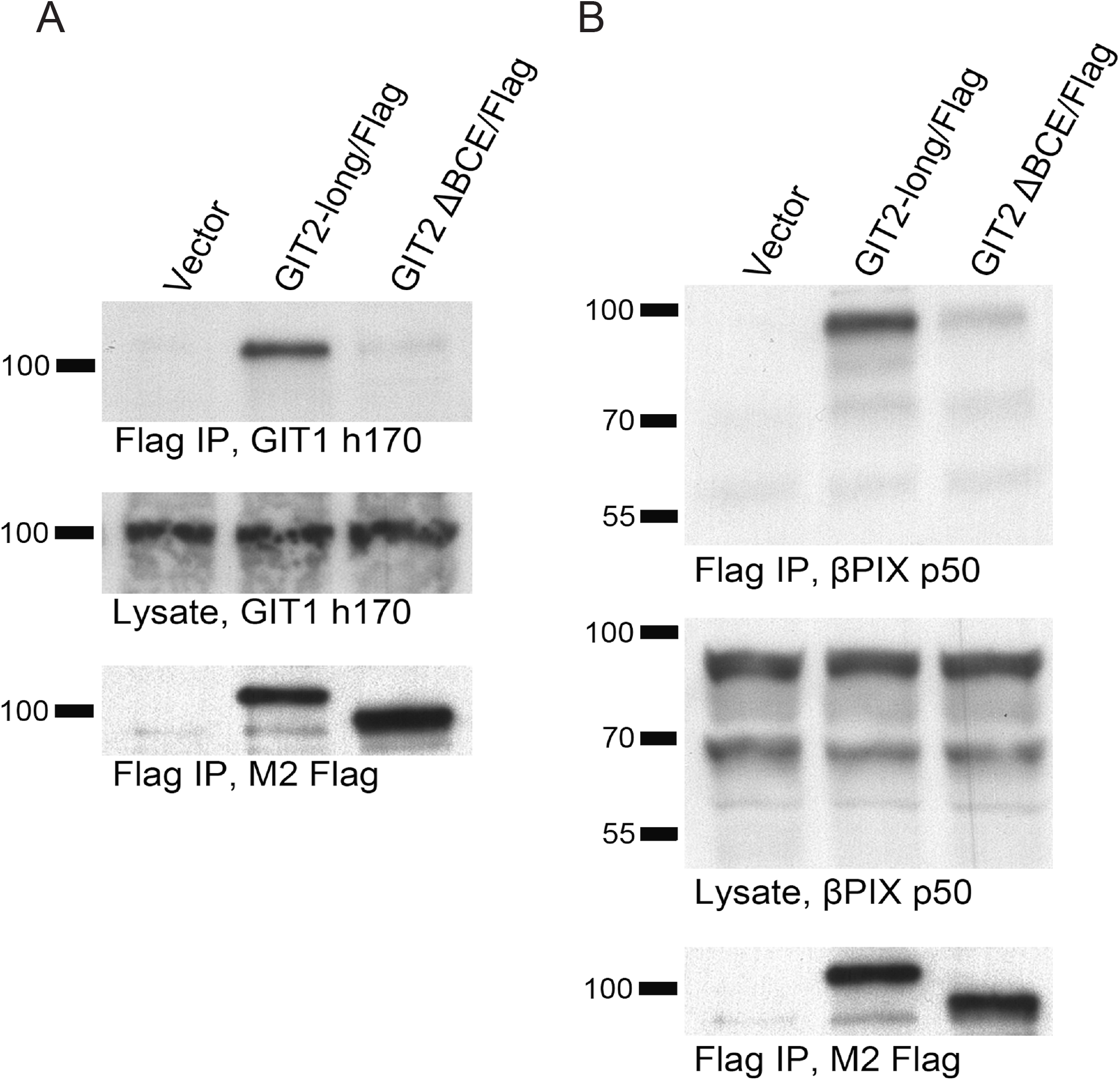
GIT2(ΔBCE) fails to dimerize or to associate tightly with PIX proteins. A) The GIT2-long/Flag or GIT2(ΔBCE)/Flag constructs were transfected into HEK293 cells, and dimerization with endogenous GIT1 was assessed following Flag immunoprecipitation. While GIT1 is associated with GIT2-long, it does not co-immunoprecipitate with GIT2(ΔBCE). B) GIT2-long/Flag or GIT2(ΔBCE)/Flag were transfected into HEK293 cells, and association with endogenous β-PIX was assessed following Flag immunoprecipitation. While β-PIX is associated with GIT2-long, it only weakly co-immunoprecipitates with GIT2(ΔBCE). Blots shown are representative of at least three independent experiments.

Because GIT protein dimerization plays an important role in forming the multimeric GIT/PIX complex (Premont, Perry et al. 2004), we also assessed the ability of GIT2(ΔBCE) to tightly associate with PIX proteins in GIT/PIX complexes (Fig 10B). COS7 cells express several β-PIX variants, and these native proteins abundantly co-immunoprecipitated with GIT2-long after forming GIT2/ß-PIX complexes. However, GIT2(ΔBCE) only weakly co-immunoprecipitated native β-PIX, consistent with reduced ability to form oligomeric GIT/PIX complexes. Instead, there is a very low but significant level of β-PIX associated with GIT2(ΔBCE) over background, consistent with weak binding of β-PIX solely to the intact Spa2 domain of the GIT2(ΔBCE) monomer, as we have seen previously for a GIT1 mutant lacking the coiled-coil dimerization domain (Premont, Perry et al. 2004).

## Discussion

In biochemical assays, GIT1 and GIT2 have generally appeared to be interchangeable (Premont, Claing et al. 2000, Vitale, Patton et al. 2000). In the brain, GIT1 and GIT2 appear to be present in nearly all neurons (Schmalzigaug, Phee et al. 2007). Since the two GIT proteins can readily heterodimerize as well as homodimerize in model cells (Premont, Perry et al. 2004), they have been presumed to be functionally redundant. However, a few studies have suggested distinct functions or distinct regulation of GIT1 and GIT2 (Brown, Cary et al. 2005, Frank, Adelstein et al. 2006, Schmalzigaug, Garron et al. 2007).

With the behavioral analysis of GIT2-deficient mice presented here, it is now very clear that GIT2-KO and GIT1-KO mice exhibit very distinct learning and memory phenotypes: GIT1-KO are markedly deficient in learning, while GIT2-KO are grossly normal. This is the first demonstration of learning and memory function in the absence of GIT2.

We find no support for the hypothesis that loss of GIT2 might lead to an ADHD-like phenotype, as has been reported for GIT1. The linkage of GIT1 to ADHD has become controversial, as we are unable to demonstrate ADHD-like behavior in GIT1-knockout mice in our tests (Martyn, Toth et al. 2018), and additional human association studies have not found an association between ADHD and GIT1 polymorphisms in several human populations (Salatino-Oliveira, Genro et al. 2012, Klein, van der Voet et al. 2015). Here we show that GIT2-KO exhibit normal spontaneous locomotor activity rather than basal hyperactivity, and fail to show evidence of amphetamine-induced locomotor suppression. However, GIT2-KO mice do not respond normally to amphetamine, since they exhibit increased sensitivity to this drug, responding to a low dose that does not activate locomotion in wildtype littermates. In contrast, loss of GIT1 leads to reduced sensitivity to amphetamine in our hands (Martyn, Toth et al. 2018). Recent reports that GIT proteins affect presynaptic neurotransmitter release (Podufall, Tian et al. 2014, Montesinos, Dong et al. 2015) confirms that GIT proteins have both presynaptic and postsynaptic roles, so further study will be required to understand how GIT2 and GIT1 differentially affect amphetamine sensitivity.

In our original characterization of GIT2, the extensive tissue-specific alternative splicing of this transcript was examined functionally by comparing only the GIT2-long and GIT2-short variants (Premont, Claing et al. 2000). Both of these forms were capable of binding strongly to PIX proteins to form GIT2/PIX complexes. However, they differed in the ability to bind to paxillin, as this interaction is mediated by the carboxyl terminal focal adhesion-targeting domain (Schmalzigaug, Garron et al. 2007) that is absent in GIT2-short, which has a truncated carboxyl terminus lacking the alternative D and E exons as well as the conserved FAT domain. From the present work, it is clear that this initial comparison was insufficient, since it did not assess the effects of loss of the of alternatively-spliced regions A, B and C. Since that time, no other studies have compared GIT2 variants, and the effects of alternative splicing have been ignored. In particular, the lack of avid, specific GIT2 antisera has hampered efforts to more carefully examine tissue-specific splicing, although it is clear that native GIT2 comes in multiple forms (Schmalzigaug, Rodriguiz et al. 2009). Further effort is needed to clarify potentially distinct roles of other GIT2 splice variants.

Interestingly, studies from the Cerione lab have all used their mouse GIT2 clone, called “CAT2”, which is a ΔBC variant (Bagrodia, Bailey et al. 1999). Because neither these workers nor anyone else has ever directly compared this GIT2(ΔBC) to any GIT2 form containing the BC region, the assumption has been made that the PIX interaction they measured using “CAT2” was as robust as that reported by other groups using GIT2 forms containing the BC region. Now it seems likely that their reported interaction was only the very weak association of a GIT2 monomer with PIX (as seen Fig 10B). In the absence of this comparison, the Cerione group created models for how PIX is activated by GIT2(ΔBC) assuming association and dissociation of monomeric GIT2 and PIX (Feng, Baird et al. 2004, Baird, Feng et al. 2005) that seemed to make little sense in the context of tightly-associated oligomeric GIT/PIX complexes, but which are clearly possible with weak monomeric association with GIT2(ΔBC) or GIT2(ΔBCE). Further work is clearly needed here to compare how weak association of monomeric GIT2(ΔBC) or GIT2(ΔBCE) with PIX may lead to PIX activation, versus how PIX is activated within stable oligomeric GIT/PIX complexes (which, it is now clear, has never been tested directly). Similarly, we predict that any GIT2 variants, in any tissue, lacking the B or C regions may similarly be unable to assemble into GIT dimers or further into oligomeric GIT/PIX complexes.

These results also highlight a common misunderstanding in the literature about the association of GIT proteins with PIX proteins. The PIX binding site on GIT1 was mapped by the Manser lab to the Spa2 repeats using a recombinant protein overlay technique that excluded oligomeric binding (Zhao, Manser et al. 2000). Our own work using co-immunoprecipitation of complexes from cells indicated that loss of the Spa2 repeats from GIT1, either through point mutations or complete deletion, is insufficient to prevent GIT association with PIX, and that in cells, GIT protein coiled-coil dimeric interactions are critical to assembling GIT/PIX complexes (Premont, Perry et al. 2004). For GIT1, and all GIT2 variants containing the complete coiled-coil region (+BC), we predict that these proteins will exist primarily if not exclusively within stable oligomeric GIT/PIX complexes, while GIT2 forms lacking the BC region will be primarily monomeric and only loosely and transiently associated with PIX trimers. It remains unknown what amount of cellular PIX is found as free PIX trimer, unassociated with GIT/PIX complexes, but the very substantial loss of brain PIX protein in the absence of GIT1 (Won, Mah et al. 2011)(Fig 8) but not in the absence of GIT2 (that is, mainly GIT2(ΔBCE)) suggests that in the brain most PIX is found within multimeric GIT1/PIX complexes with only a minor fraction that is only loosely bound to monomeric GIT2(ΔBCE) and other variants like it. Instead, the small amount of PIX interaction with GIT2(ΔBCE) is consistent with loose association of one GIT2(ΔBCE) with PIX solely through Spa2 interactions. Additionally, β-PIX in particular also exists as multiple splice variants, one class of which lacks the coiled-coil region responsible for PIX trimerization (Kim, Kim et al. 2000, Koh, Manser et al. 2001). It is thus tempting to speculate that dimerization-deficient GIT2 monomers associate primarily with trimerization-deficient β-PIX forms in a loose, regulated association of two monomers, in contrast to the apparently constitutive GIT/PIX oligomers assembled from GIT and PIX protein variants containing functional coiled-coil domains.

There are several functional consequences of brain GIT2(ΔBCE) not forming oligomeric GIT2/PIX complexes. In the absence of GIT2 expression, the native GIT1 is able to maintain normal levels of PIX proteins in the brain through stabilization within GIT1/PIX complexes. That is, mice lacking GIT2 have near-normal GIT/PIX complex levels and localization, which are able to scaffold PAK appropriately within synapses. Other partners requiring intact GIT/PIX complexes for proper localization and function (Zhou, Li et al. 2016) also should be relatively unaffected by loss of GIT2 in the brain. The inability to properly localize PAK function in the absence of GIT1 leads to abnormal hippocampal synaptic structural plasticity (Martyn, Toth et al. 2018) and presumably also to defective long-term potentiation, and thus poor learning and memory behavior; whereas all these functions are normal (or are expected to be) in GIT2-deficient mice.

On the other hand, hippocampal neuron dendritic spine density is reduced similarly by loss of either GIT2 (shown here) or GIT1 (Menon, Deane et al. 2010, Martyn, Toth et al. 2018), suggesting that this role of GIT proteins does not require scaffolding or crosstalk within GIT/PIX complexes. A report suggesting that the ArfGAP function of GIT1 is important for regulating spine stability (Rocca, Amici et al. 2013) hints that GIT2 might also regulate spines in an Arf-dependent manner, independent of PIX or of PIX partners such as PAK.

Overall, we conclude that GIT2(ΔBCE), a prominent GIT2 isoform in the hippocampus and amygdala and throughout the entire brain, is unable to dimerize or participate in oligomeric GIT/PIX complexes, and therefore cannot affect PIX-dependent pathways (particularly PAK pathways) that appear to be critical for supporting learning and memory function. Thus global loss of GIT2, and particularly of this brain GIT2(ΔBCE) variant, does not lead to noticeable loss of brain PIX protein due to destabilization, nor to the severe learning deficits observed in GIT1-KO or a-PIX-KO mice.

## Acknowledgements

We thank Pamela Bonner for mouse husbandry and genotyping, Shivani Salunke for assistance with spine density measurements, Rick Cerione (Cornell University, Ithaca, New York) for the p50 β-PIX antiserum, and Hisatake Sabe (Hokkaido University, Sapporo, Japan) and Yuichi Mazaki (Kumamoto University, Kumamoto, Japan) for the NEO replacement GIT2-knockout mouse line.

Supported by NIH R21 MH090556 and Department of Defense Medical Research and Development Program (DMRDP) contract W81XWH-11-2-0112 to RTP. The funders had no role in study design, data collection and analysis, decision to publish, or preparation of the manuscript.

Author contributions
RS, RTP, conceived and planned project; KT, ACM, NB, WK, UA, RS, RMR, RTP, performed experiments; KT, ACM, RS, RMR, RTP, analyzed data; KT, ACM, RS, RTP, wrote manuscript; WCW, edited manuscript; RTP, obtained funding.

